# Rapid neurostimulation at micron scale with optically controlled thermal-capture technique

**DOI:** 10.1101/2024.07.15.603492

**Authors:** Alexey M. Romshin, Nikolay A. Aseyev, Olga S. Idzhilova, Alena A. Koryagina, Vadim E. Zeeb, I. I. Vlasov, Pavel M. Balaban

## Abstract

Precise control of cellular temperature at the microscale is crucial for developing novel neurostimulation techniques. Here, we study the effect of local heat on the electrophysiological properties of cells at the subcellular level using a cutting-edge micrometer-scale thermal probe, the diamond heater-thermometer (DHT). Experiments on primary neuronal cultures and HEK293 cells revealed that millisecond heat pulses could induce reversible changes in membrane potential and elicit ionic displacement currents. At local temperatures close to 50 °C, a rapid increase in cellular response by an order of magnitude was observed, attributed to local phase changes in the phospholipid membrane at the point of contact with the DHT. This allows the cell membrane to be effectively and reproducibly captured by temperature, referred to as thermal-capture mode (TCM). Once transition to TCM occurred, even lower temperatures (<35 °C) elicited depolarization up to 10 mV in neurons, sufficient for triggering action potentials with rates up to 30 Hz. Additionally, the impact of high temperatures beyond the physiological range on the electrophysiology of the cell was assessed. These findings enhance the understanding of how local heat affects cellular functions and provide insights into the thermal modulation of cell activity.

## Introduction

Thermal variations at microscale can affect electrophysiological parameters of living cells, enabling the control of membrane potential and the selective stimulation of single cells both *in vitro* [1], [2], [3], [4], [5] and *in vivo* [3], [6], [7], [8].

The mechanisms underlying the dependence of electrical parameters of the cell on temperature have been extensively studied in recent years. A number of approaches aimed to deliver a precise amount of heat in order to control the electrophysiological state of living cells have been developed. One of the most elaborated and wide-used, is a local pulsed infrared stimulation, exploiting the absorption of water to rapidly increase temperature at the cellular level [6], [9]. It was shown that even slight local heating triggers the changes in membrane potential and overall permeability of the cell membrane. An infrared-induced thermal effect results in reversible changes in the electrical capacitance of the plasma membrane of cells [1], depolarizing the cells and generating action potentials. Subsequently, Beier et al. [10] proposed that infrared radiation causes structural changes in the cell membrane, leading to the formation of transient nanopores which increase the non-selective ionic permeability of the membrane, contributing to cell depolarization. This effect was also reversible, allowing the membrane to return to its original state 20 minutes after the cessation of infrared stimulation, indicating a mechanism that could be used repeatedly for therapeutic purposes without permanent damage to the cells. Another family of methodologies employed photoabsorption materials, such as metallic [2] and semiconductor [4] nanoparticles, as well as biopolymers [11], for effective light-to-heat conversion. It was shown that plasmonic nanoparticles can be conjugated to high-avidity ligands having an affinity for specific proteins on the neuronal membrane in primary culture [2]. It was revealed that the efficacy of the neuronal response depends on the target, being higher in the case of TRPV1 and voltage-gated sodium channels, and less when the ligand targets the ATP-gated P2X3 receptor. Nonetheless, changes in membrane capacitance and depolarization currents were observed in the targeted neurons regardless of the type of ion channel. Therefore, the action of local heating on the cell appears to be more dependent on the thermal membrane properties rather than mediated solely by ion channels-dependent temperature perception.

Despite the clarity of previous studies, the mechanism of the membrane potential shift due to the effect of local heat on cellular compartments and the cell as a whole remains incompletely determined. Partly, constraints arise from optical techniques, as both infrared and photothermal materials make it impossible to affect the electrophysiological processes with high spatial resolution and are limited to the scale of the whole cell soma (∼10 µm). Although plasmonics typically employs metallic nanoparticles of several tens of nanometers for thermal stimulation, conjugating a multitude of these nanoparticles can result in the entire cell surface being exposed to heat. Moreover, aforementioned methods either required preliminary temperature calibration using external thermometers [1], [4], [10], sometimes giving inaccurate temperatures, especially for nanoscale heat sources [2], or were applied blindly, so that just the relative changes in the optical power applied to the heater were determined [12]. Such approaches do not provide an adequate assessment of the amount of heat delivered to the biological system, leading to incorrect interpretation of the conditions, under which cellular processes are initiated.

In this study, the effect of local heat on the electrophysiological properties of cells at the subcellular level was investigated using a state-of-the-art thermal microprobe, known as the diamond heater-thermometer (DHT), which combines the functions of a thermometer and a heater in a single micrometer scale particle. Experimentally, DHT was capable of controlling local temperature with an accuracy <0.2 °C in the vicinity of the cell’s phospholipid bilayer, occupying less than 3% of its surface. Millisecond local heat pulses induced reversible changes in membrane potential and elicited capacitive currents in cultured neurons and HEK293 cells. At local temperatures close to 50 °C, a rapid increase in cellular response by an order of magnitude was observed, which we attribute to local phase changes in the phospholipid membrane at the point of contact with DHT, allowing the cell to be effectively and reproducibly captured by temperature in the so-called thermal capture mode (TCM). Once transition to TCM occurred, even lower temperatures (<35 °C) elicited depolarization up to 10 mV in neurons sufficient for triggering the action potentials (APs) at the rate up to 30 Hz. In addition, the impact of high temperatures far from the physiological range on the electrophysiology of the cell was estimated with the focus on the AP generation. Present findings enhance an established understanding of how local heat affects cellular functions and provide valuable insights into thermal modulation of cell activity.

### Experimental section

Experimental measurements were carried out on custom-built optical setup (**Fig. 1a**), combining the possibilities of potential fixation on the neuronal membrane (whole-cell patch clamp) and fully optical temperature control using DHT. The base of the setup was an optical microscope with the ability of translational movement along three Cartesian coordinates. The imaging system of living cells was based on the use of DGC contrast (see Methods) in combination with the Olympus 40× water immersion objective (NA=1.0). Micromanipulators with high spatial accuracy of movement were used for precision positioning of the patch electrode headstage and the DHT. The design of the diamond thermometer heater is based on the use of a submicron glass capillary and a single fluorescent diamond nanoparticle, slightly melted to the tip of the capillary (**Fig. 1b**, for DHT design details see [13], [14]). To excite the fluorescence of the SiV centers, a laser source emitting at a wavelength of 640 nm was used, equipped with TTL-input which was employed for generation of heat pulses with arbitrary duration (from 200 ns) and repetition frequency (up to 10 MHz). Spectrometer Ocean Insight QE Pro (1800 mm^-1^, slit 100 μm) was employed to register SiV fluorescence spectra and subsequently determine the position of the ZPL using a self-designed algorithm described in [13]. In most experiments involving micro-scale thermal impact on living cells, a diamond heater-thermometer of 1.4 μm diameter (DHT1.4) was used (Fig. 1b (i)). Only in specifically noted cases smaller particles of approximately 500 nm were used (DHT0.5, **Fig. 1b (ii)**).

**Fig. 1.**
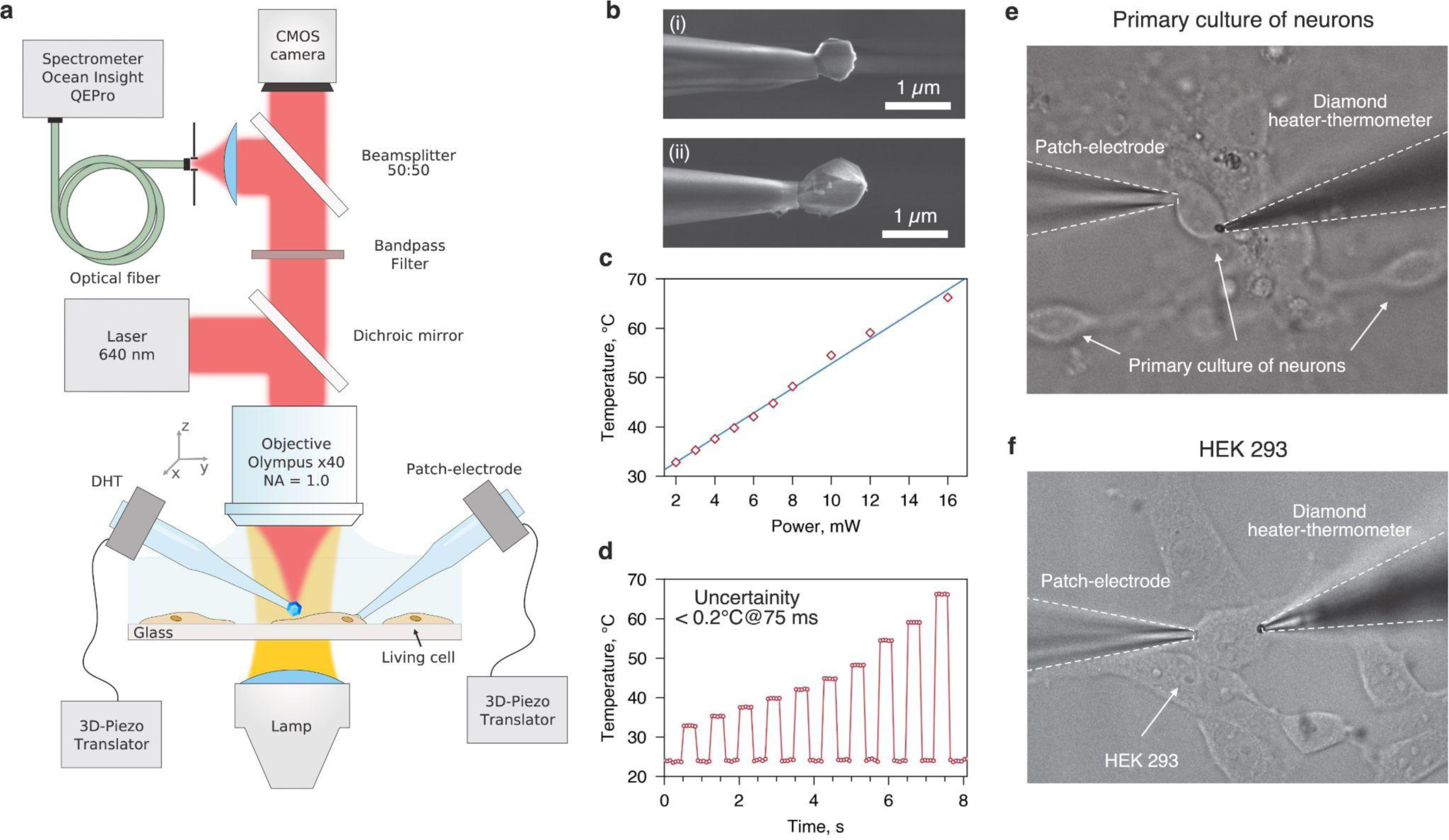
(**a**) A schematic illustration of an experimental setup tailored to electrophysiological and thermometric measurements at the same time. (**b**) SEM-image of DHTs with 500 nm (**i**) and 1.4 μm (**ii**) diamonds’ sizes. (**c**) Power dependence of DHT local temperature response in saline solution maintained under ambient conditions at 23.1 °C. Slope coefficient responsible for the ability of DHT to heat local volume of extracellular solution is 2.5 ± 0.1 °C/mW. (**d**) Time course of the DHT-temperature measured with 75 ms acquisition at a variety of laser powers. S.e.m. is less than 0.2 °C at each power step. Dependencies (**c**)-(**d**) were obtained for 1.4 μm DHT. (**e**) Optical DGC-image of the primary culture of neurons, patch electrode and DHT arrangement. (**f**) Optical DGC-image of HEK 293 cells, patch electrode and DHT arrangement.

## Results

At the beginning, for DHT1.4, the temperature of the diamond particle, which was in local thermodynamic equilibrium with the extracellular solution, was measured as a function of the power of the laser radiation exciting the fluorescence (**Fig. 1c**). The range of achievable ΔT was broad, from a few degrees up to 60°C, with an increase in power by 1 mW leading to a local temperature change on the surface of the diamond by +2.5°C. An instance of a time course of measured temperature with a time step of 75 ms under laser pulses (225 ms) with various powers is presented in the **Fig. 1d**, from where the detection accuracy of <0.2 °C is clearly determined. Note that due to the high thermal conductivity of diamond, the temperature distribution within it is quasi-uniform, making the measured temperature value consistent throughout the diamond particle. Further heat propagation into the water-based extracellular medium decreases rapidly on submicron scales, in good agreement with Fourier’s law [13]. An indicated spatial scales correspond to equilibrium times that are short compared to macroscopic systems. It was theoretically shown in [15] that the characteristic time τ_c_ for the cooling of a spherical particle in the liquid medium is determined by equation 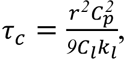, where *r* and *C*_p_ are radius and heat capacity per volume of the particle, *k*_&_ and *C*_&_ are liquid thermal conductivity and heat capacity per volume, respectively. Сonsequently, 1 μm diamond particle in aqueous medium reaches equilibrium with its surroundings in no more than 1 μs. Additionally, a recent study [16] experimentally measured the thermal relaxation times of gold nanoparticles (50 nm fraction) in water, finding them to be 200-300 ps. According to these results and discussion in [17], we assume that the influence of the temperature gradient establishment processes might be neglected in comparison to the physiological time scale: electrophysiological response to micro-/nanoscale heat is always postponed in time, while for IR-laser heating (10-50 μm^3^) thermal relaxation takes at least several tens of milliseconds [1], [10].

Electrophysiological patch clamp measurements were conducted in the primary culture of neurons or HEK293 cells in the whole-cell configuration. To effectively expose the cell to local heat, the DHT1.4 was positioned as close as possible to the selected cell with minimal mechanical damage. Specifically, the DHT1.4 was placed a few micrometers above the cell, then brought closer until a slight characteristic bend of the membrane indicated contact. The absence of current through the cell membrane as the DHT1.4 approached ruled out the activation of mechanosensitive channels. At the same time, the DHT1.4 was positioned laterally away from the controlling patch electrode to avoid non-physiological current changes due to thermal alteration of the patch micropipette resistance. Although control measurements did not show significant thermal changes in current through the patch electrode in the absence of a living cell (**Fig. S1**), the DHT1.4’s position was chosen to be far from the patch electrode (as depicted in **Fig. 1e-f**) to avoid possible heat-related artifacts. Also, application of laser pulse alone in the absence of DHT at focal point did not produce any electrophysiological response (**Fig. S2**).

As expected from previous studies with infrared and plasmonic thermal stimulation [1], [2], in current clamp mode, the neurons were depolarized from their resting potential with heat pulses produced by the DHT1.4 (**Fig. 2a**). It can be logically assumed that the DHT1.4 heater, occupying no more than 3% of the cell soma area, could only lead to minor potential changes. However, we revealed two contrasting modes of local temperature impact on neurons, presumably differing in the degree of the DHT1.4’s interaction with the cell membrane and the phase state changes of the latter. In the first mode, the DHT1.4 slightly touched the cell membrane without altering its phase properties, operating in probe-mode (P-mode, PM). The depolarization amplitude increased with the rise in peak heating temperature, reaching a maximum of 1.1 mV at remarkably high values of 70-80 °C. Surprisingly, even after such thermal exposures, neurons remained physiologically stable (able to generate action potentials and maintain resting potential) for at least 30 minutes in gap-free protocols (**Fig. S3**), indicating probable low thermal toxicity with short-duration temperature pulses at micro- and nanoscale. The typical depolarization rise time in P-mode was weakly dependent on the heating amplitude and ranged from 0.5 to 1 ms.

**Fig. 2.**
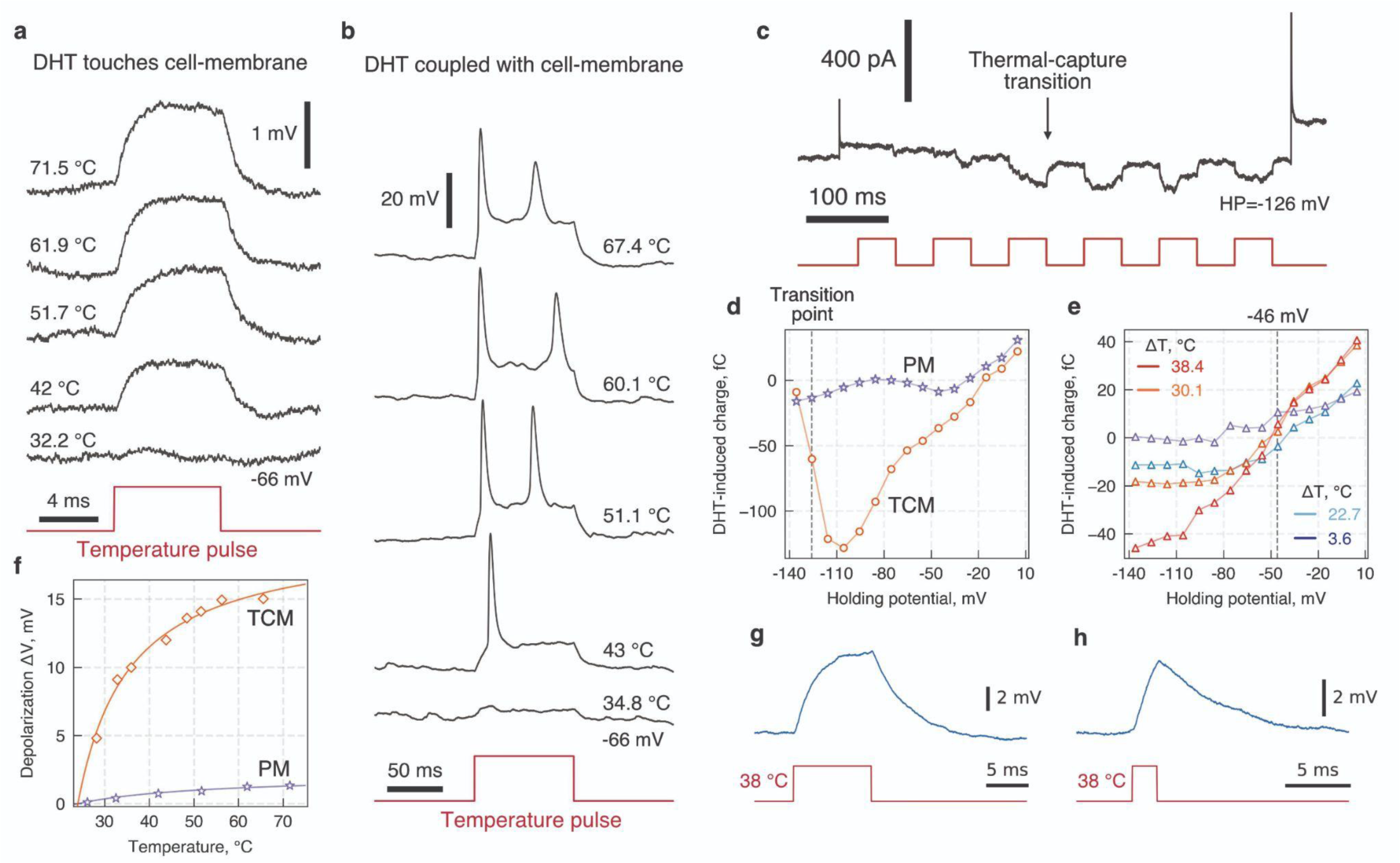
Thermally-induced changes of electrophysiological properties in cultured neurons. (a)-(b) Reversible depolarization dynamics of current-clamped neurons under local heating protocols recorded in two different modes of DHT-to-membrane affinity: (a) probe mode (PM, DHT gently touches the cell membrane) and (b) thermal-capture mode (TCM, DHT strongly coupled with cell membrane). The red line shows the heat pulse shape with a duration of 10 ms. (c) Currents recorded in voltage-clamped neurons held at -126 mV evoked by 10 Hz train of 55 °C heat pulses. At the third heat pulse where the current changes breach, a transition between PM and TCM is observed. (d) A heat-induced charge transfer through the cell membrane with respect to holding voltage (Q-V curves) obtained for PM and TCM operating modes. N=6 for each measurement. (e) Q-V response of the voltage-clamped neuron to the application of heat pulse with different amplitudes ΔT in TCM: 3.6 °C (dark blue), 22.7 °C (cyan), 30.1 °C (orange) and 38.4 °C (red). (f) Maximum induced changes of membrane potential with respect to the absolute DHT temperature in PM (purple) and TCM (orange) modes. (g)-(h) A depolarization response of a current-clamped neuron to the 10 ms (g) and 2 ms (h) heat pulses with an absolute temperature amplitude of 38 °C in TCM. Red lines show the heat pulses timing, cyan lines reflect voltage changes.

For the majority of cells, we revealed a more pronounced mode of potential change in the local temperature field, occurring at an energy threshold of 50-55 °C and heat pulse duration of ∼50 ms (**Fig. 2b**). Beyond these threshold conditions, the depolarization amplitude elicited by the same temperature stimulus increased 15-fold up to 16 mV, while the time required for saturation of the depolarization voltages increased to 3-4 ms, depending on the cell soma size and contact area. Even with a lower thermal stimulus at 37 °C, the depolarization response was around 10 mV (**Fig. 2g**). Fig. 2c shows the voltage-clamp time course of the current under a 10 Hz sequence of 50-ms heat pulses (T_heat_ = 55 °C). Initial pulses elicited only slight depolarizing currents; however, by the third pulse, the energy barrier was gradually breached, and from that moment, the peak current values increased dramatically. Subsequent protocols of local heating with varying durations and amplitudes always led to reversible depolarization over hundreds of trials in both current-clamp and voltage-clamp conditions. Presumably, the DHT1.4 could lead to local melting of the membrane and establishment of a closer contact with it — transitioning to a thermal-capture mode (TC-mode, TCM). In fact, along with the report in [1] on the infrared light-induced local discoloration of *X. laevis* oocytes, we also observed the similar effect on the neuronal surface after removing the thermal probe and sometimes stretched filaments of the membrane entrained with the melted membrane part. It should be noted that after the DHT1.4 was completely removed from the extracellular solution, the life cycle of the neurons continued, they fully responded to incoming synaptic signals, and were capable of demonstrating both spontaneous and evoked electrophysiological activity.

The transition from PM to TCM is accompanied by a sharp nonlinear increase of inward currents and, subsequently, a corresponding charge-voltage (Q-V) response to thermal stimuli at holding voltage -126 mV and T_heat_ = 55 °C (**Fig. 2d**). After the TC-mode has been established, the charge increases linearly with voltage, with a slope coefficient of 1.44±0.12 fC/mV, reaching the reversal at -36 mV for this cell. The average reversal potential across N=5 neurons is -46 mV, which statistically shows little dependence on local temperature magnitude (**Fig. 2e**). The observed reversal potential variation of ±15 mV is likely due to the increasing contribution of activated ion channels as the holding current rises along with probable change in surface charge density in the close proximity of the cell membrane [18]. In TCM, the charge flowing through the membrane increases with temperature, peaking at ΔT between 35 and 50 °C. Further increases in heating magnitude on the millisecond timescale compromise cell integrity, leading to loss of resting potential and electrophysiological dysfunction.

For a quantitative comparison of P-mode and TC-mode, the maximal depolarizing ΔV at various local temperature amplitudes was measured (**Fig. 2f**). Saturation behavior with increasing temperature was clearly observed for both modes and was well described by the equation *ΔV* = *ΔV*_∞_ ⋅ (*T* − *T_sol_*)/(*T* − *T_sol_* + Δ*T*_sat_), where *ΔV_∞_* is the high-temperature depolarization limit, Δ*T*_sat)*_is the saturation relative temperature corresponding to the curve inflection, *T*_sol_ is the temperature of the ambient extracellular solution. Based on the fitting values in TCM, the saturation limit *ΔV_∞_^TCM^* = 19.7 mV was reached at *ΔV_sat_^TCM^* = 11 °C, while the corresponding parameter in PM is almost 20 times less, with a depolarizing limit *ΔV_∞_^PM^* = 2.1 mV and a saturation temperature *ΔV_sat_^PM^* = 29.7 °C.

Like the external electrical stimuli, the thermal pulses in TC-mode led to generation of action potentials in neurons upon reaching the threshold depolarization values. Since in current-clamp mode, the voltage response to local temperature is inert, the depolarization magnitude can be adjusted not only by the amplitude of the thermal pulse but also by its duration. Note that DHT1.4 was capable of inducing depolarizatiove currents with a frequency at least 2 kHz and a pulse duration of 100 µs (**Fig. S4**). **Fig. 3a** shows a comparison of the voltage time courses, whilst 10 Hz sequences of pulses of different durations were applied, as the DHT1.4 was heated to 40 °C. Starting with 5 ms, thermal depolarization reaches the threshold potential with a 20% probability, while pulses with a duration of 10 ms reliably lead to action potential generation. The detailed overlay of 10 ms pulses on the voltage time course is shown in **Fig. 3b**, demonstrating accurate “on-demand” generation of action potentials, at the thermal pulse rate of 10 Hz.

**Fig. 3.**
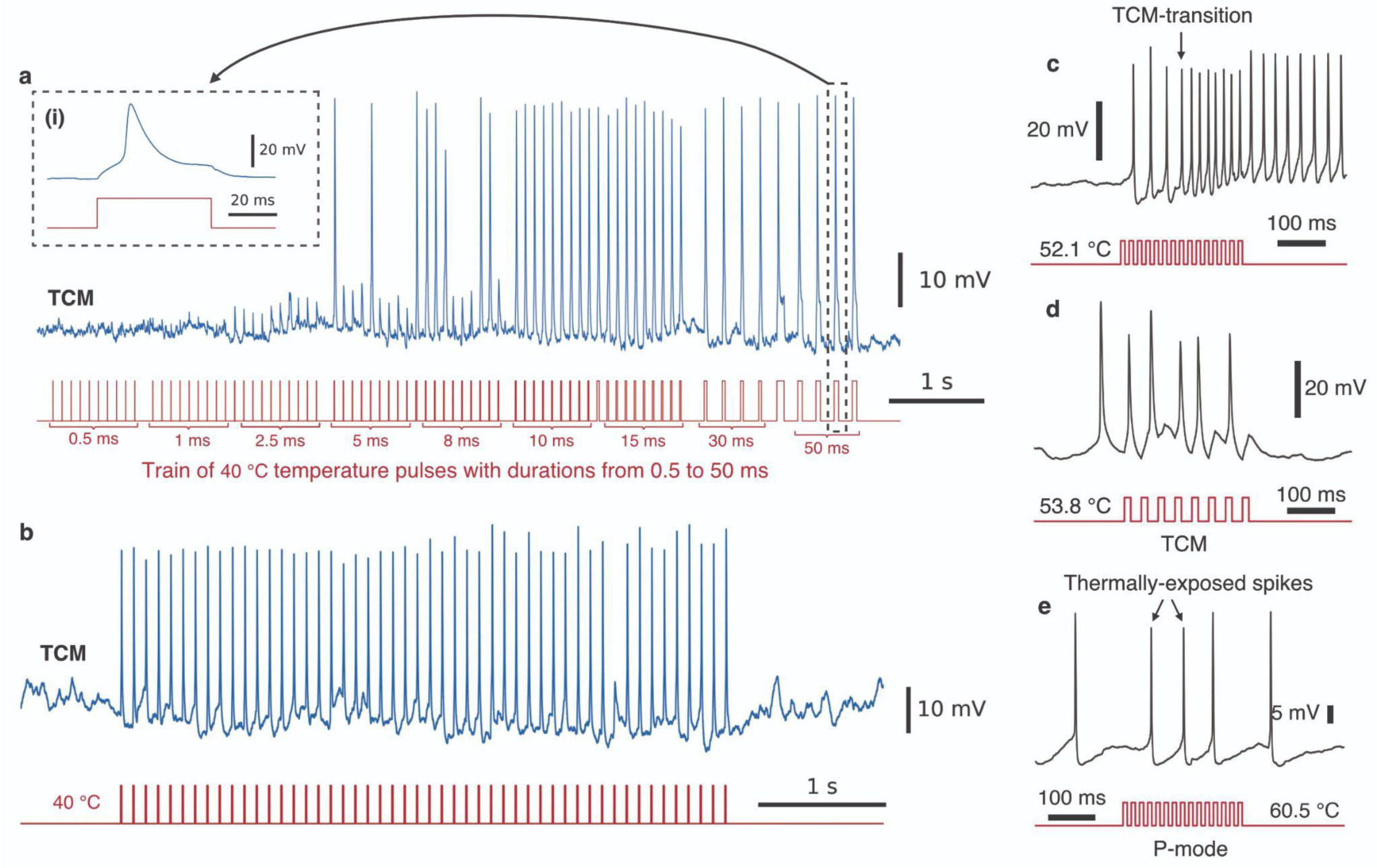
TCM-induced generation of action potentials in cultured neurons at a variety of heat pulse durations and amplitudes. (a) Train of heat pulses (T_heat_ = 40 °C) with durations of 0.5 ms, 1 ms, 2.5 ms, 5 ms, 8 ms, 10 ms, 15 ms, 30 ms, 50 ms in chronological order. An inset shows zoomed action potential waveform elicited by the 50 ms pulse. (b) A 10 Hz train of 8 ms heat pulses reliably initiates action potentials. (c) 60 Hz train of 8 ms thermal pulses and a current-clamp transition to TCM. Tightly packed action potentials in the second part of the burst are thermally evoked in TC-mode. (d) Initiation of action potentials in TCM: 30 Hz train, 10 ms duration. (e) P-mode: 60 Hz train of 8 ms thermal pulses; responses superimposed with spontaneous electrical activity of neurons. Two action potentials exposed to heat (marked with arrows) are decreased in amplitude by ∼10%.

The most suitable frequency for thermal-induced action potential generation in most neurons was found to be 10 Hz. However, some cells were able to respond at frequencies up to 30 Hz (**Fig. 3c**) and 25 Hz (**Fig. 3d**), where dense overlapping of action potentials during the repolarization and auto-hyperpolarization phases is observed, as well as a slight (10-15%) decrease in their amplitude. Notably, a similar relative decrease in the AP peak amplitude occurring during heat pulses was also observed in P-mode (**Fig. 3f**). We measured the action potential durations during electrical stimulation in the absence of thermal impact and found that the action potential duration at 90% repolarization (APD90) in the most cases was 25-30 ms. Therefore, rate of the thermal stimulation is naturally limited to 30-40 Hz due to the prolonged APD in cultured neurons [19] and, in part, due to related factors such as the refractory period of the preсeding action potential overlap with the subsequent thermal depolarization. Additionally, the thermal excitability was studied with a smaller size of the heat source (DHT0.5). It was found that application of a 5 Hz train of heat pulses with ΔT=26 °C also evokes APs in primary culture of neurons (**Fig. S5**).

An observation of amplitude decrease of APs in both TC-mode and P-mode led us to consider the waveform of APs in detail. In order to understand how the local temperature affects the peak amplitude and duration of APs, an electrical stimulation and thermal exposure were combined. Specifically, a short-term electrical stimulus was necessary for the controlled depolarization of the cell to the threshold potential, while a local thermal pulse of the same duration played a controlling role by altering the shape of the evoked action potential (AP). Fig. 4a-b illustrates the evolution of action potential shape with increasing temperature in the case of strong coupling between the DHT1.4 and cell membrane (TCM) for both current- and voltage-clamp modes, respectively. As in previous studies on AP properties modulation with macroscale temperature [20], the spike amplitude decreased as the local T_heat_ increased (**Fig. 4c**). This effect was most clearly manifested in the current-clamp mode, where spike amplitudes dropped by up to 30% (from 60 to 42 mV) at T_heat_ ∼ 57 °C. In contrast, action potentials recorded in the voltage-clamp mode experienced only a slight decrease in amplitude of 2-3%, which can be partly explained by the fact that current is less susceptible to possible changes in cell capacitance than voltage. Similar to the peak amplitude, a trend towards acceleration of the depolarization and repolarization phases of APs was observed (**Fig. 4d**). The full width at half maximum (FWHM or APD50) decreased in both current-clamp and voltage-clamp modes by 33% and 25%, respectively. As one might expect, action potential rise and decay times followed the trend of the FWHM and decreased with temperature. Yet, the depolarization time reproducibly decreased with the temperature rise, becoming 2-4% faster than the repolarization time.

**Fig. 4.**
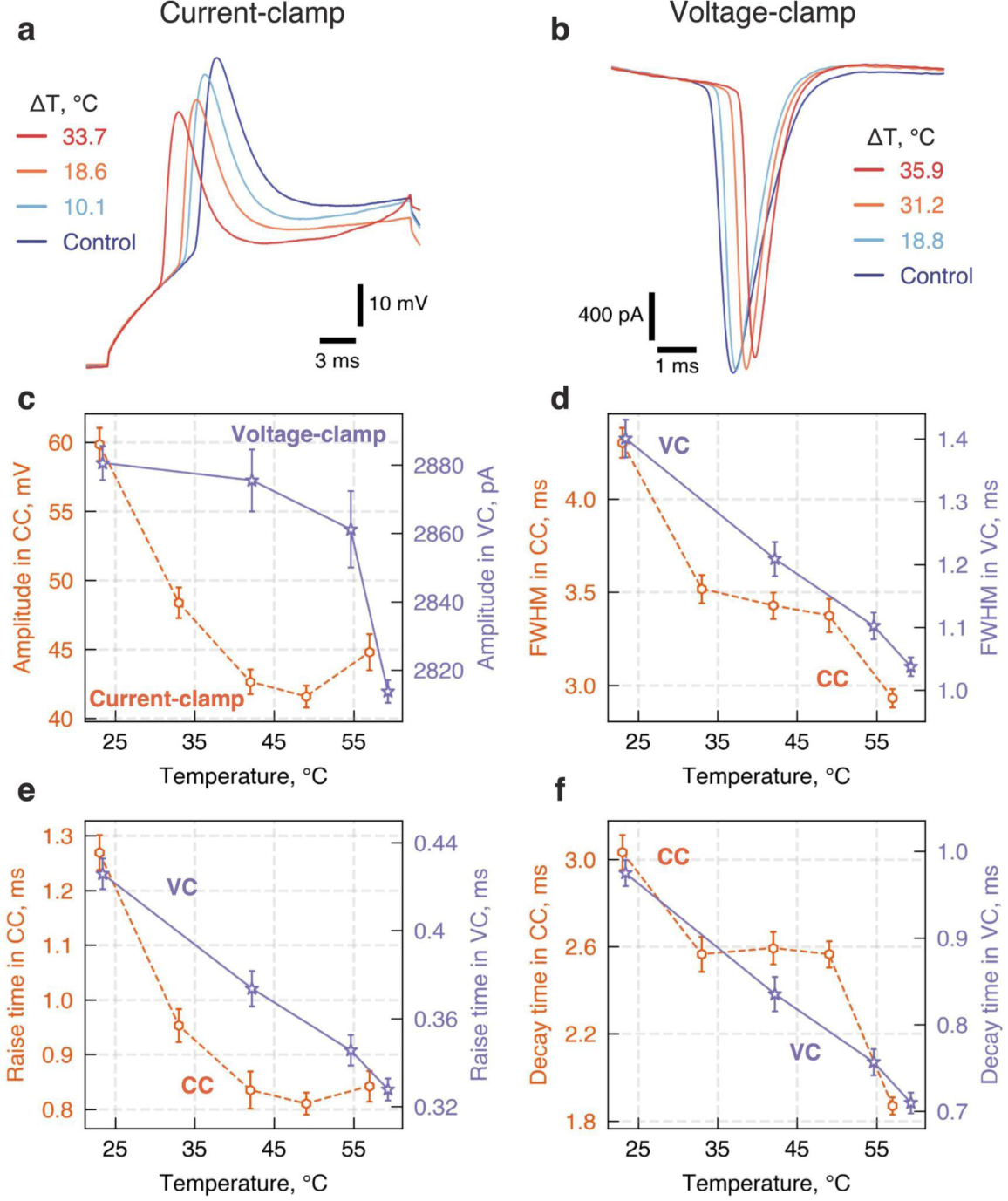
Local temperature changes the kinetic parameters of primary culture neurons’ APs. (a) Voltage response of current-clamped neuron to the application of 200 pA depolarizing current for 30 ms during 30 ms thermal pulses in TCM: control (blue), ΔT = 10.1 °C (cyan), ΔT = 18.6 °C (orange) and ΔT = 33.7 °C (red). (b) Current response of voltage-clamped neuron to the application of 20 mV depolarizing voltage for 15 ms during 15 ms thermal pulses in TCM: control (blue), ΔT = 18.8 °C (cyan), ΔT = 31.2 °C (orange) and ΔT = 35.9 °C (red). (c-f) Dependence of parameters of electrically evoked action potentials on the applied T_heat_ for both voltage-clamp and current-clamp modes: peak amplitude (с), full width at half maximum (APD50) (d), rise time (e) and decay time (f). The data were acquired in both voltage-clamp (purple stars, solid lines) and current-clamp (orange hexagons, dashed lines) modes. N=20 for each experimental point. Error bars stand for standard error of the mean (s.e.m.). The lines are presented for better visibility.

The above results obtained on cultured neurons demonstrate the reliable capability of thermal control over the potential and current of excitable cells. To determine the possible impact of the local temperature upsurge on non-excitable cells and examine whether the TCM transition is an intrinsic property exclusive to excitable cells, we performed the DHT1.4 stimulation measurements in whole-cell-clamped HEK293 cells. Initially, a 500 ms 42.7 °C thermal pulse was applied to the cell three times (**Fig. 5b**) to stimulate the TCM transition. While the first pulse elicited a slight inward current of ∼10 pA, the second one enhanced the response up to 40 pA by the end of the thermal exposure. The third pulse did not result in a significant increase in depolarization currents, leading us to conclude that TC-mode had been achieved. Subsequently, the application of a 50 ms temperature pulse at T_heat_ = 55 °C produced the same current response as observed in neurons (**Fig. 5a**), with an inward direction at high negative holding potentials and outward at positive potentials. In contrast to neurons, the reversal of the current sign for HEKs occurred at a higher holding potential, -26 mV.

**Fig. 5.**
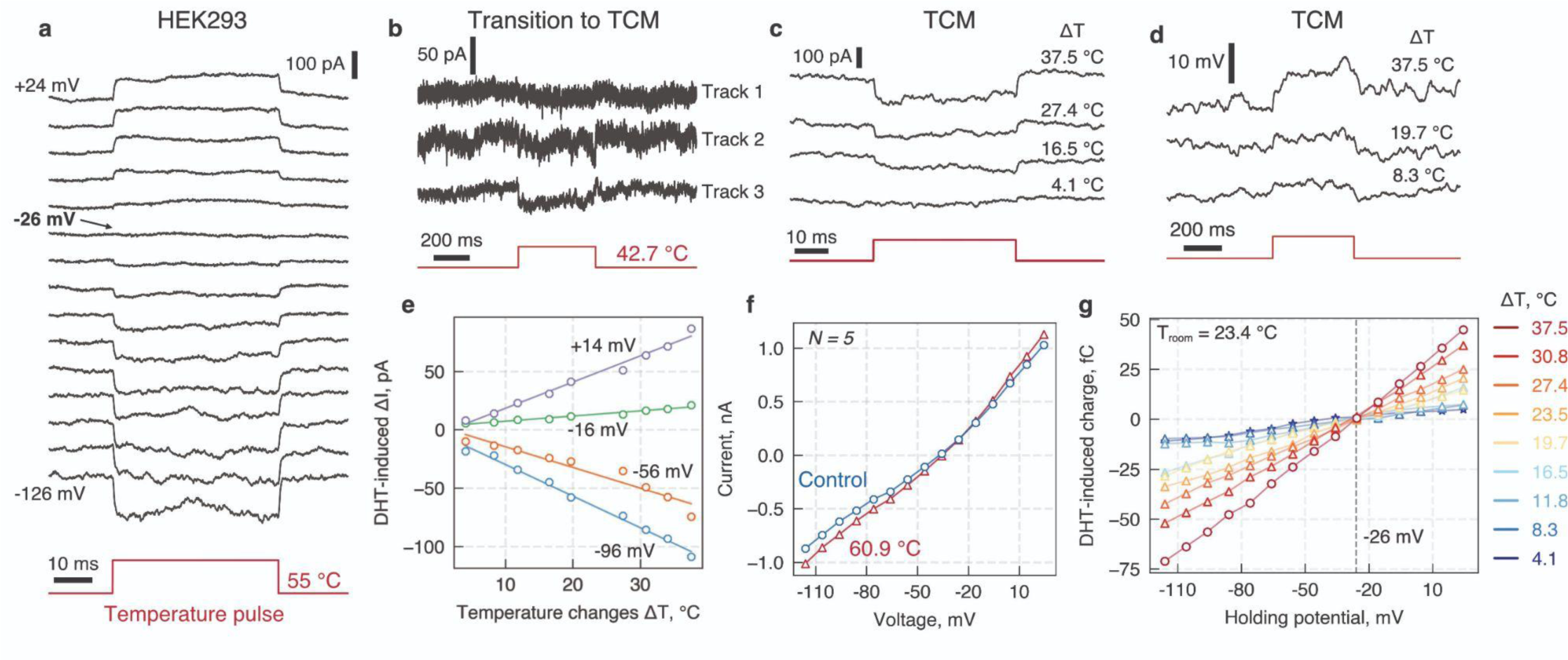
DHT1.4 thermal pulses elicit current and potential changes in HEK293. (a) Current- voltage response to the application of 50 ms 55 °C pulses. An arrow at -26 mV trace indicates the reversing of the current. (b) 500 ms 42.7 °C pulse leads to transition from P-mode to TC- mode. (c) Current response in TCM to 50 ms heat pulses of 4.1, 16.5, 27.4, 37.5 °C at holding potential -66 mV. (d) Voltage response in TCM to 500 ms heat pulses of 8.3, 19.7, 37.5 °C. (e) A comparison of DHT1.4-induced current changes to local temperature pulses at -96 mV (blue), -56 mV (orange), -16 mV (green) and +14 mV (purple). (f) Volt-ampere characteristics acquired for HEK cells: control (blue) and exposed to 50 ms 60.9 °C pulses (red). N=5 for each measurement. (g) Q-V response of the voltage-clamped HEK293 to the application of a variety of ΔT: 4.1, 8.3, 11.8, 15.5, 19.7, 23.5, 27.4, 30.8, 37.5 °C.

The dynamics of heat-induced currents in HEK293 cells was studied at different local temperatures. The amplitude of the current response exhibited linear growth with increasing local temperature, showing no signs of saturation even at ΔT > 35 °C (**Fig. 5e**). The transition time required to reach the new current level was found to be independent of temperature, and approximately 1 ms. This time constant was used to determine the magnitude of the flowing charge and to construct the corresponding Q-V curves (**Fig. 5g**). The capacitive charge (-75 pA at -126 mV and ΔT = 37.5 °C) was comparable in magnitude to the corresponding parameter for neurons. Contrary to data obtained in neurons, analysis of 5 HEK cells showed no divergence in reversal potential; at different local temperatures, the current sign change always occurred at -26 mV.

Finally, the response of membrane resistance and capacitance to local heat in TCM for different types of cells was evaluated (**Fig. 6**). For this, a train of 200 ms 10 mV depolarizing voltage pulses was applied to the cell. By analyzing the renewed level of current and calculating the charge over transient current response, these parameters were estimated. As expected, the capacitance was found to increase by 5-6% under elevation of local temperature to 40 °C, while the resistance synchronously decreased from 10% up to 40%. However, since HEK 293 form syncytium of cells and primary neuronal cultures produce the extracellular matrix, a relatively high native capacitance of the cells was observed.

**Fig. 6.**
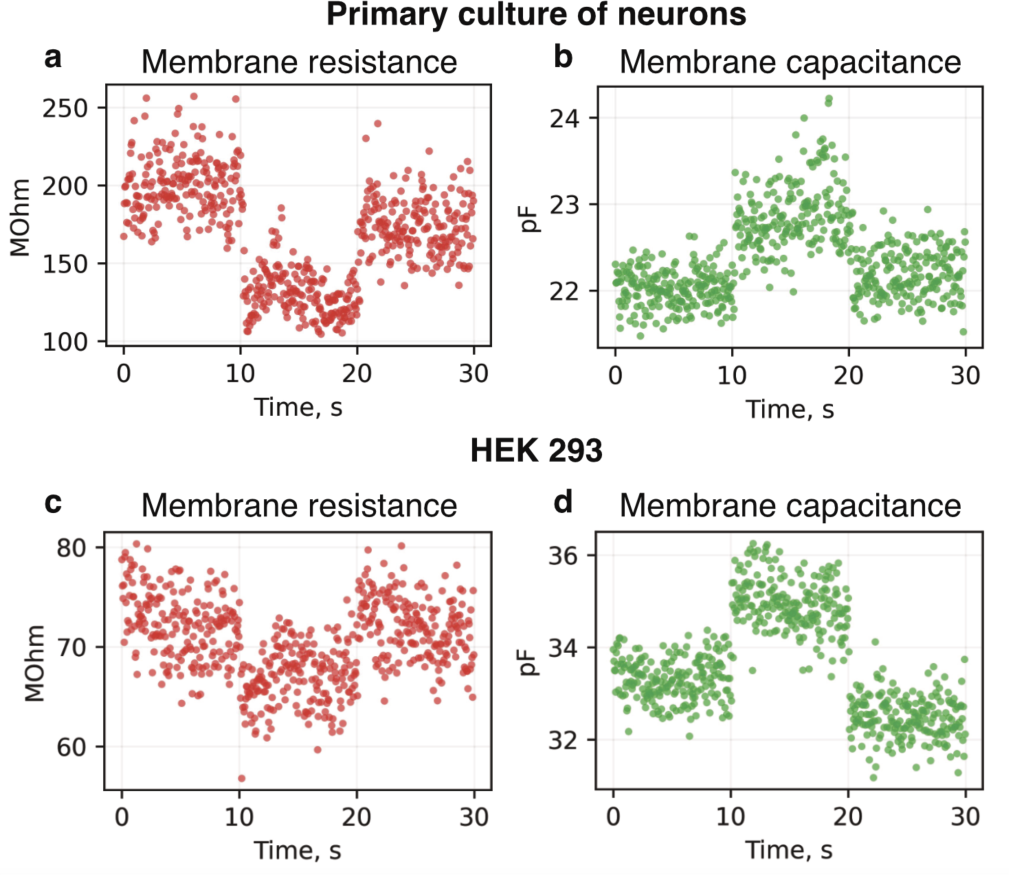
Alteration of membrane resistance (a, c) and capacitance (b, d) of the cultured neurons and HEK cells by 10 s 40 °C DHT1.4 heating (from 10 s to 20 s).

## Discussion

We demonstrated the impact of local heat on the electrophysiological properties of cells at the subcellular level using an advanced thermal microprobe. Two distinct modes of action of local heat on a living cell: probe-mode (PM) and thermal-capture mode (TCM), were clearly observed. In PM, where the DHT touches the cell membrane, local heating reproducibly altered the membrane potential and elicited depolarization currents within the millisecond timescale required to reach the renewed current level. A typical voltage shift for local temperatures ranging from 25 to 70 °C is approximately 1 mV. In TCM, it was found that the application of a 50 °C temperature step induces an order-of-magnitude greater increase in cellular response, allowing the cell to be captured by temperature and depolarized by up to 20 mV. This abrupt and reversible transformation in response suggests a critical threshold temperature at which the phospholipid membrane undergoes local phase changes, which lead to both increased ionic permeability and depolarization currents, and impaired cellular susceptibility to temperature.

Observations of this transition in both cultured neurons and HEK293 cells suggest that local temperature stimulation affects their membranes in a similar manner, indicating a common underlying mechanism of TCM transition in excitable and non-excitable cells.

In previously reported studies, predominantly the entire surface of the cell was exposed to heat, leading to a displacement of the membrane potential by amounts ranging from several millivolts to 30 mV for infrared and plasmonic thermal stimulation, respectively [1], [2]. Given that the DHT occupies only ∼3% of the cell soma surface, the equivalent depolarization for the entire cell in PM is around 30 mV, which is consistent with previous studies [1], [10]. Meanwhile, a similar estimation for TCM at a local temperature of 37 °C gives ∼330 mV, significantly exceeding the acceptable physiological limit. Therefore, heating at the cellular scale in earlier experiments would lead to irreversible damage at these temperatures rather than triggering phase changes in the phospholipid membrane bilayer as the DHT does.

Generally, heat affects the properties of the phospholipid bilayer by altering its fluidity and permeability. At higher temperatures, phospholipid molecules gain kinetic energy. The resulting increased fluidity can disrupt the membrane structure producing the ’melting’ effect, as the phospholipids move further apart. Thus the membrane permeability is increased, potentially allowing normally impenetrant molecules to pass through more easily (local poration). An observed increase in cell capacitance by 5-6% in TCM alters the ionic environment in the vicinity of the phospholipid bilayer, eliciting depolarization currents through the cell membrane. According to the recently proposed MechanoElectrical Thermal Activation (META) theory [18], the total displacement current is contributed by capacitive currents and concurrent changes of the surface charge densities in Stern layers, with corresponding thermal capacitance increase rate of ∼0.3%/°C. At the same time, the decrease of cell membrane resistance and increase of inward currents potentially refer to either reversible changes in joint contribution of ion channels, particularly sodium (Na+) and potassium (K+) [21], [22], or the possible formation of local pores in the lipid bilayer with enhanced fluidity [10], [23], facilitating non-selective ion transport. Eventually, local heat above physiological range may cause denaturation of proteins on the cell membrane [24], leading to the dysfunction of specific cellular receptors. However, since the voltage response to DHT stimuli in TCM is reversible over time, we believe this thermodynamic route is less probable.

The abovementioned factors partly explain changes in excitability and AP shape in neurons at local temperature elevation. First, a faster kinetics of ion channels may lead to a quicker inactivation of Na+ channels and a more rapid activation of K+ channels, reducing the overlap between inward Na+ currents and outward K+ currents [20], [22]. The rapid inactivation of Na+ channels results in a shorter duration of the action potential, as the period during which Na+ ions enter the neuron is curtailed. Consequently, the action potential peak amplitude diminishes since fewer Na+ ions contribute to the depolarization phase. Second, an increased membrane fluidity may facilitate the inactivation of Na+ channels and the activation of K+ channels, thereby supporting the rapid repolarization of the action potential and contributing to a reduced AP amplitude and duration.

In conclusion, we demonstrate the capability of spatiotemporally confined thermal stimuli to significantly alter the physiological properties of cells. A novel approach based on interaction between plasma membrane and local heat sources is promising for controlling cellular electrophysiological states, although temperatures beyond physiological limit pose risks of thermal damage, necessitating careful calibration and control to ensure cell viability and function. The ability to trigger action potentials at sub-physiological temperatures immediately after the thermal-capture transition along with reduction of APDs, opens uncharted avenues for refined neurostimulation techniques.

## Acknowledgements

The work was supported by the grant of the Russian Science Foundation No. 23-14-00129 (https://rscf.ru/project/23-14-00129/). Authors express gratitude to Prof. Evgeny Nikitin (Laboratory of Cellular Neurobiology of Learning, IHNA RAS) for meaningful comments on the work and I.V. Smirnov (Laboratory of Cellular Neurobiology of Learning, IHNA RAS) for technical help. The Institute of Higher Nervous Activity and Neurophysiology thanks Wonder Technologies LLC for providing the DHT setup to conduct the above studies.

## Methods

All experimental procedures were in compliance with the Guide for the Care and Use of Laboratory Animals published by the National Institute of Health and were approved by the Ethical Committee of the Institute of Higher Nervous Activity and Neurophysiology, Russian Academy of Sciences.

### Electrophysiology

Whole-cell recordings with patch electrodes were obtained from cultured primary neurons and HEK293TN cells using Dodt-Gradient-Contrast System (DGC, Luigs & Neumann, Ratingen, Germany) optics and videomicroscopy mounted on LNScope microscope (Luigs & Neumann, Ratingen, Germany) with 40× water immersion objective. For the precise positioning of the patch and DHT-holding pipettes, the rig was equipped with motorized micromanipulators Мini 25 (Luigs & Neumann, Ratingen, Germany) mounted on an air table. The bath was filled with HBSS solution (Hank’s Balanced Salt Solution with glucose and HEPES: 10 mM glucose, 10 mM HEPES, 139 mM NaCl, 5.3 mM KCl, 1.26 mM CaCl_2_, 0.5 mM MgCl_2_, 4.16 mM NaHCO_3_, 0.34 mM Na_2_HPO_4_, 0.44 mM KH_2_PO_4_, 0.4 mM MgSO_4,_ pH 7.4). The patch electrodes were filled with a potassium gluconate based solution (140 mM potassium gluconate, 5 mM KCl, 4 mM Mg-ATP, 0.3 mM Na_2_-GTP, 10 mM sodium phosphocreatine, 10 mM HEPES) and had a resistance of 4–6 MΩ for neurons and 6–9 MΩ for HEK293TN cells. Recordings were made with a ELC-03XS amplifier (NPI Electronics, Germany) in the current clamp or voltage clamp mode. After amplification, the data were digitized at 50 kHz and fed into a computer using Digidata 1550B interface and Clampex ver. 10.7 software (both from Molecular Devices, USA). All the data obtained was corrected to liquid junction potential (LJP) which is +16.1 mV for solutions used in this work.

### Culturing cells

Primary neuronal cultures were prepared by the common technique [25], from the cortices of ICR mouse pups on postnatal day 0-1. Cortical tissue was extracted in ice-cold Dulbecco’s Modified Eagle Medium (DMEM, Thermo Fisher Scientific, USA) with 15 mM HEPES, minced and transferred to a tube with prewarmed 0.08% solution of trypsin in DMEM. After 15 minutes of incubation at 37°C, the tube was centrifuged. The precipitate was washed with DMEM to get rid of the enzyme, and then dissolved in Neurobasal™-A medium, supplemented with B27™ (50X) (both, Thermo Fisher Scientific) and Alanyl-glutamine (200 mM, PanEko, Russia). Tube contents were triturated several times, and after each trituration, a fraction of single cell suspension was transferred to another tube. Thus obtained, cell suspension was rediluted in NBM and plated on Poly-L-Lysine-coated 10 mm glass coverslips. Cultures were kept in a CO2 incubator at 5% CO2, 37°C, 95% humidity. 1/3 of the culture medium was replaced with fresh NBM once every 3-4 days.

HEK293TN cells were grown in Dulbecco modified eagle medium supplemented with 10% fetal bovine serum. One or two days before patching, cells were plated on a 4-well plate (d=15.4 mm) with an average density 65 cells/mm^2^.

## Supplementary Information

### 1. Patch-electrode voltage under local heating protocols

**Fig. S1.**
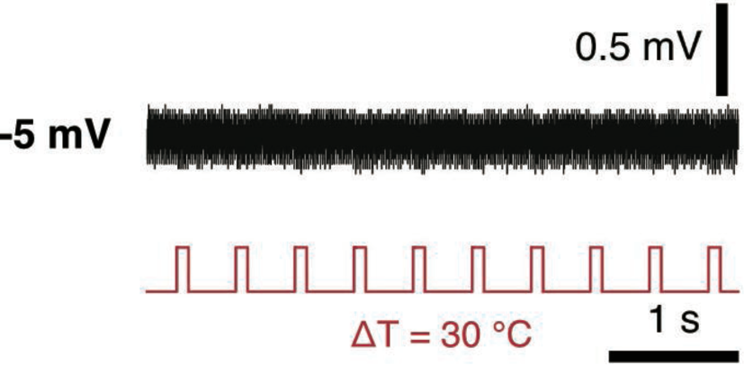
Changes in the glass electrode voltage under a 2 Hz train of 100 ms heat pulses (ΔT_heat_ = 30 °C). The distance between electrode and DHT was ∼1 µm.

### 2. Influence of 640 nm laser irradiation on electrophysiological properties of cells

**Fig. S2.**
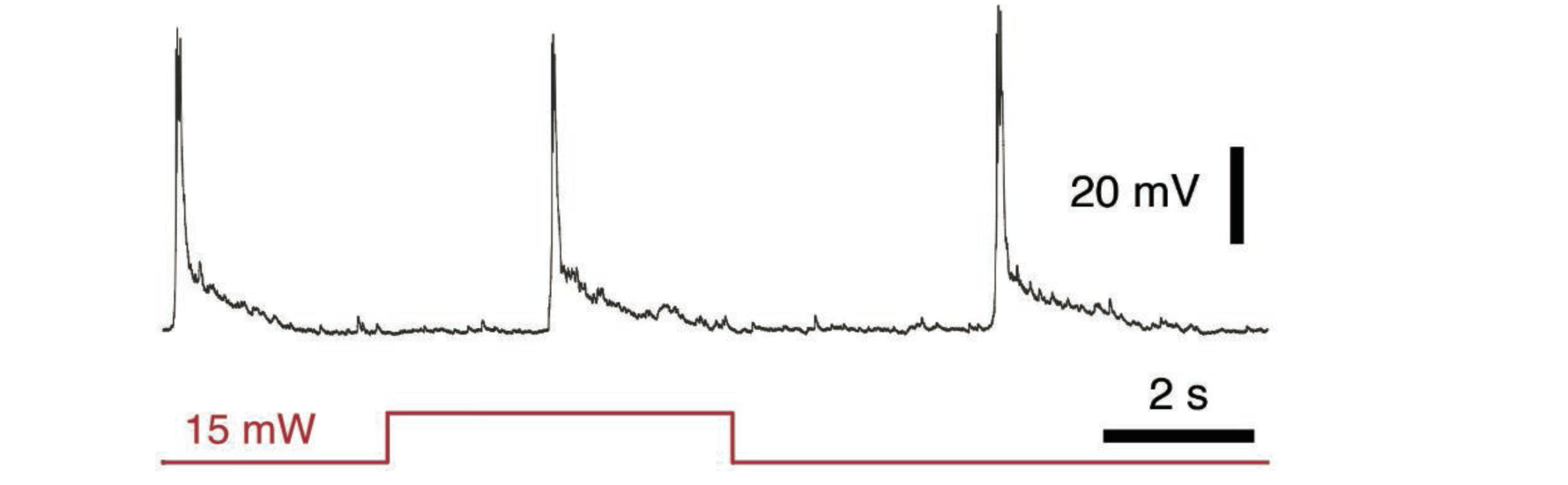
Application of 15 mW 5s laser pulse alone in the absence of DHT at focal point did not produce any electrophysiological response in neurons.

### 3. Resistance of neurons to non-physiological temperature at microscale

**Fig. S3.**
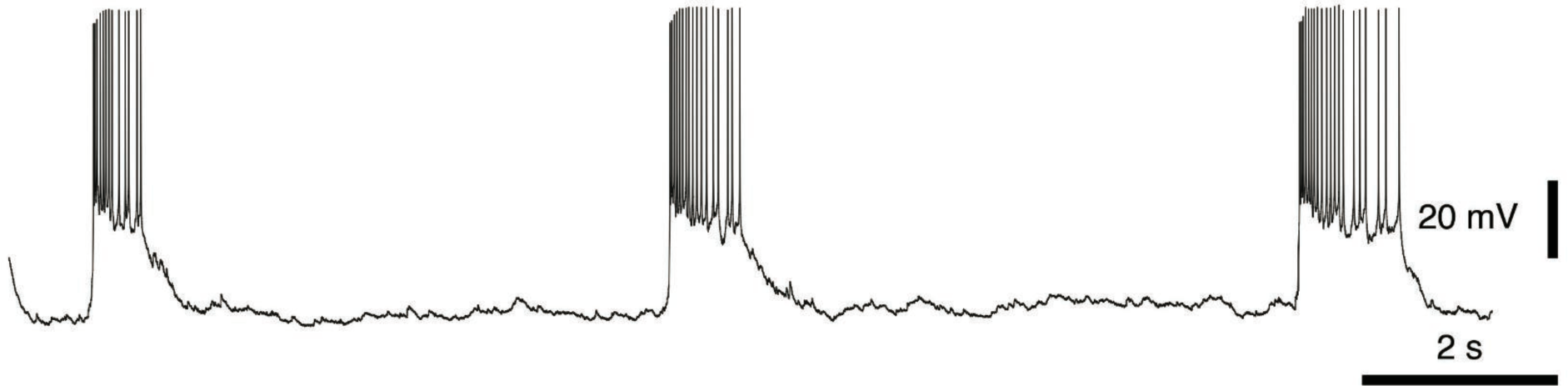
Membrane potential and burst activity in a current-clamped neuron 30 min after cessation of the sequence of ten heat pulses with duration 1 ms and T_heat_ = 80 °C.

### 4. Depolarization currents in neurons with short thermal pulses

**Fig. S4.**
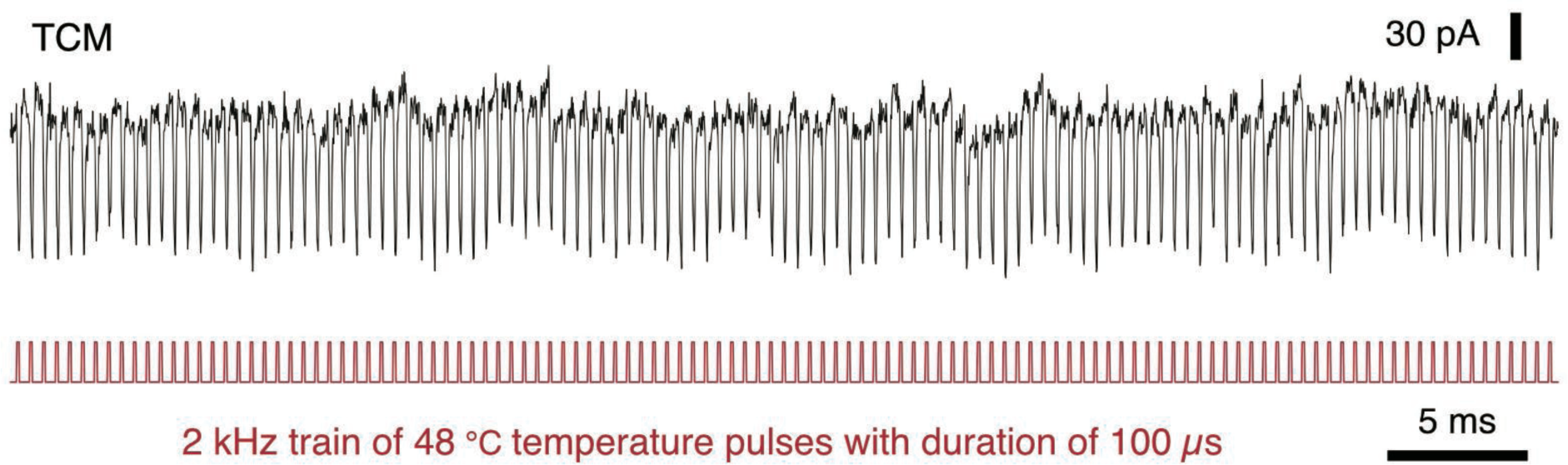
TCM-induced depolarization currents in cultured neurons at a 2 kHz train of 100 µs heat pulses (T_heat_ = 48 °C) produced by DHT1.4.

### 5. Thermal excitability of neurons with DHT0.5

**Fig. S5.**
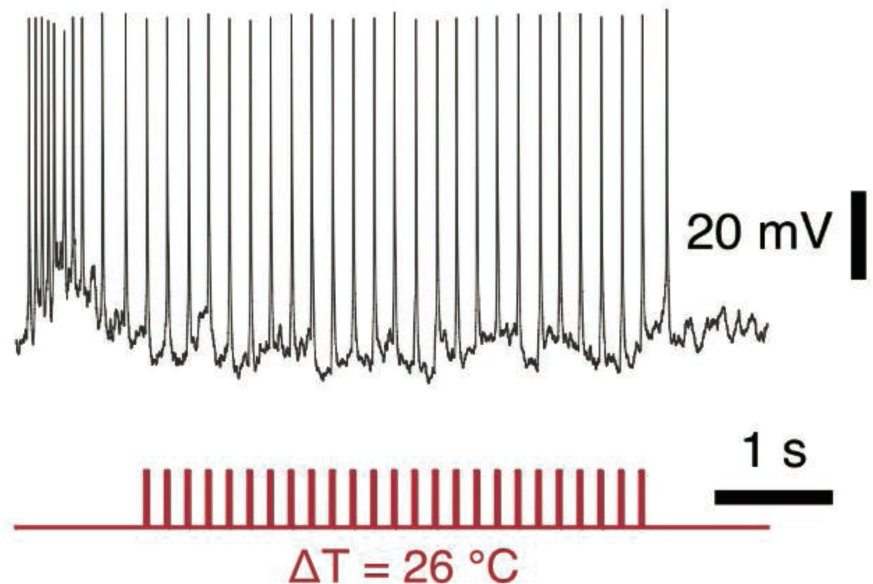
TCM-induced generation of action potentials in cultured neurons at a 5 Hz train of 25 ms heat pulses (ΔT_heat_ = 26 °C) produced by DHT of 500 nm in size.

## References

[1] M. G. Shapiro, K. Homma, S. Villarreal, C.-P. Richter, and F. Bezanilla, ‘Infrared light excites cells by changing their electrical capacitance’, Nat. Commun., vol. 3, no. 1, p. 736, Mar. 2012, doi: 10.1038/ncomms1742.

[2] J. L. Carvalho-de-Souza, J. S. Treger, B. Dang, S. B. H. Kent, D. R. Pepperberg, and F. Bezanilla, ‘Photosensitivity of Neurons Enabled by Cell-Targeted Gold Nanoparticles’, Neuron, vol. 86, no. 1, pp. 207–217, Apr. 2015, doi: 10.1016/j.neuron.2015.02.033.

[3] Y. Jiang et al., ‘Neural Stimulation In Vitro and In Vivo by Photoacoustic Nanotransducers’, Matter, vol. 4, no. 2, pp. 654–674, Feb. 2021, doi: 10.1016/j.matt.2020.11.019.

[4] Y. Jiang et al., ‘Heterogeneous silicon mesostructures for lipid-supported bioelectric interfaces’, Nat. Mater., vol. 15, no. 9, pp. 1023–1030, Sep. 2016, doi: 10.1038/nmat4673.

[5] C. -P. Richter, A. I. Matic, J. D. Wells, E. D. Jansen, and J. T. Walsh, ‘Neural stimulation with optical radiation’, Laser Photonics Rev., vol. 5, no. 1, pp. 68–80, Jan. 2011, doi: 10.1002/lpor.200900044.

[6] J. Wells et al., ‘Optical stimulation of neural tissue in vivo’, Opt. Lett., vol. 30, no. 5, p. 504, Mar. 2005, doi: 10.1364/OL.30.000504.

[7] J. Wells, C. Kao, E. D. Jansen, P. Konrad, and A. Mahadevan-Jansen, ‘Application of infrared light for in vivo neural stimulation’, J. Biomed. Opt., vol. 10, no. 6, p. 064003, 2005, doi: 10.1117/1.2121772.

[8] R. Chen, G. Romero, M. G. Christiansen, A. Mohr, and P. Anikeeva, ‘Wireless magnetothermal deep brain stimulation’, Science, vol. 347, no. 6229, pp. 1477–1480, Mar. 2015, doi: 10.1126/science.1261821.

[9] A. D. Izzo et al., ‘Laser Stimulation of Auditory Neurons: Effect of Shorter Pulse Duration and Penetration Depth’, Biophys. J., vol. 94, no. 8, pp. 3159–3166, Apr. 2008, doi: 10.1529/biophysj.107.117150.

[10] H. T. Beier, G. P. Tolstykh, J. D. Musick, R. J. Thomas, and B. L. Ibey, ‘Plasma membrane nanoporation as a possible mechanism behind infrared excitation of cells’, J. Neural Eng., vol. 11, no. 6, p. 066006, Dec. 2014, doi: 10.1088/1741-2560/11/6/066006.

[11] N. Martino et al., ‘Photothermal cellular stimulation in functional bio-polymer interfaces’, Sci. Rep., vol. 5, no. 1, p. 8911, Mar. 2015, doi: 10.1038/srep08911.

[12] P. Urban, S. R. Kirchner, C. Mühlbauer, T. Lohmüller, and J. Feldmann, ‘Reversible control of current across lipid membranes by local heating’, Sci. Rep., vol. 6, no. 1, p. 22686, Mar. 2016, doi: 10.1038/srep22686.

[13] A. M. Romshin et al., ‘A new approach to precise mapping of local temperature fields in submicrometer aqueous volumes’, Sci. Rep., vol. 11, no. 1, p. 14228, Dec. 2021, doi: 10.1038/s41598-021-93374-7.

[14] A. M. Romshin, V. Zeeb, E. Glushkov, A. Radenovic, A. G. Sinogeikin, and I. I. Vlasov, ‘Nanoscale thermal control of a single living cell enabled by diamond heater-thermometer’, Sci. Rep., vol. 13, no. 1, p. 8546, May 2023, doi: 10.1038/s41598-023-35141-4.

[15] O. M. Wilson, X. Hu, D. G. Cahill, and P. V. Braun, ‘Colloidal metal particles as probes of nanoscale thermal transport in fluids’, Phys. Rev. B, vol. 66, no. 22, p. 224301, Dec. 2002, doi: 10.1103/PhysRevB.66.224301.

[16] M. Hu and G. V. Hartland, ‘Heat Dissipation for Au Particles in Aqueous Solution: Relaxation Time versus Size’, J. Phys. Chem. B, vol. 106, no. 28, pp. 7029–7033, Jul. 2002, doi: 10.1021/jp020581+.

[17] V. Zeeb, M. Suzuki, and S. Ishiwata, ‘A novel method of thermal activation and temperature measurement in the microscopic region around single living cells’, J. Neurosci. Methods, vol. 139, no. 1, pp. 69–77, Oct. 2004, doi: 10.1016/j.jneumeth.2004.04.010.

[18] M. Plaksin, E. Shapira, E. Kimmel, and S. Shoham, ‘Thermal Transients Excite Neurons through Universal Intramembrane Mechanoelectrical Effects’, Phys. Rev. X, vol. 8, no. 1, p. 011043, Mar. 2018, doi: 10.1103/PhysRevX.8.011043.

[19] Z. Sun, D. J. Williams, B. Xu, and J. A. Gogos, ‘Altered function and maturation of primary cortical neurons from a 22q11.2 deletion mouse model of schizophrenia’, Transl. Psychiatry, vol. 8, no. 1, p. 85, Apr. 2018, doi: 10.1038/s41398-018-0132-8.

[20] M. Volgushev, T. R. Vidyasagar, M. Chistiakova, T. Yousef, and U. T. Eysel, ‘Membrane properties and spike generation in rat visual cortical cells during reversible cooling’, J. Physiol., vol. 522, no. 1, pp. 59–76, Jan. 2000, doi: 10.1111/j.1469-7793.2000.0059m.x.

[21] M. J. Van Hook, ‘Temperature effects on synaptic transmission and neuronal function in the visual thalamus’, PLOS ONE, vol. 15, no. 4, p. e0232451, Apr. 2020, doi: 10.1371/journal.pone.0232451.

[22] Y. Yu, A. P. Hill, and D. A. McCormick, ‘Warm Body Temperature Facilitates Energy Efficient Cortical Action Potentials’, PLoS Comput. Biol., vol. 8, no. 4, p. e1002456, Apr. 2012, doi: 10.1371/journal.pcbi.1002456.

[23] T. Heimburg, ‘Lipid ion channels’, Biophys. Chem., vol. 150, no. 1–3, pp. 2–22, Aug. 2010, doi: 10.1016/j.bpc.2010.02.018.

[24] J. R. Lepock, ‘Protein Denaturation During Heat Shock’, in Advances in Molecular and Cell Biology, vol. 19, Elsevier, 1997, pp. 223–259. doi: 10.1016/S1569-2558(08)60079-X.

[25] G. M. J. Beaudoin et al., ‘Culturing pyramidal neurons from the early postnatal mouse hippocampus and cortex’, Nat. Protoc., vol. 7, no. 9, pp. 1741–1754, Sep. 2012, doi: 10.1038/nprot.2012.099.

